# Improving the discovery of rare variants associated with alcohol problems by leveraging machine learning phenotype prediction and functional information

**DOI:** 10.1101/2023.09.11.557163

**Authors:** Mohammad Ahangari, Amanda Elswick Gentry, Mohammed F. Hassan, Tan Hoang Nguyen, Kenneth S. Kendler, Silviu-Alin Bacanu, Roseann E. Peterson, Brien P. Riley, Bradley T. Webb

**Affiliations:** Beyster Center for Psychiatric Genomics, Department of Psychiatry, University of California San Diego, La Jolla, CA, USA; Virginia Institute for Psychiatric and Behavioral Genetics, Virginia Commonwealth University, Richmond, VA, USA; Department of Psychiatry, Virginia Commonwealth University, Richmond, VA, USA; SUNY Downstate Health Sciences University, Department of Psychiatry and Behavioral Sciences, Institute for Genomics in Health, Brooklyn, NY, USA; Department of Human and Molecular Genetics, Virginia Commonwealth University, Richmond, VA, USA; GenOmics and Translational Research Center, RTI International, Research Triangle Park, NC, USA

## Abstract

Alcohol use disorder (AUD) is moderately heritable with significant social and economic impact. Genome-wide association studies (GWAS) have identified common variants associated with AUD, however, rare variant investigations have yet to achieve well-powered sample sizes. In this study, we conducted an interval-based exome-wide analysis of the Alcohol Use Disorder Identification Test Problems subscale (AUDIT-P) using both machine learning (ML) predicted risk and empirical functional weights. This research has been conducted using the UK Biobank Resource (application number 30782.) Filtering the 200k exome release to unrelated individuals of European ancestry resulted in a sample of 147,386 individuals with 51,357 observed and 96,029 unmeasured but predicted AUDIT-P for exome analysis. Sequence Kernel Association Test (SKAT/SKAT-O) was used for rare variant (Minor Allele Frequency (MAF) < 0.01) interval analyses using default and empirical weights. Empirical weights were constructed using annotations found significant by stratified LD Score Regression analysis of predicted AUDIT-P GWAS, providing prior functional weights specific to AUDIT-P. Using only samples with observed AUDIT-P yielded no significantly associated intervals. In contrast, *ADH1C* and *THRA* gene intervals were significant (False discovery rate (FDR) <0.05) using default and empirical weights in the predicted AUDIT-P sample, with the most significant association found using predicted AUDIT-P and empirical weights in the *ADH1C* gene (SKAT-O *P* _Default_= 1.06 x 10^-9^ and *P* _Empirical weight_ = 6.25 x 10^-11^). These findings provide evidence for rare variant association of the *ADH1C* gene with the AUDIT-P and highlight the successful leveraging of ML to increase effective sample size and prior empirical functional weights based on common variant GWAS data to refine and increase the statistical significance in underpowered phenotypes.

## Introduction

Alcohol use disorder (AUD) and problematic alcohol use (PAU) are moderately heritable conditions with significant social and economic impact [1, 2]. Twin-based heritability (h^2^) of AUD is estimated at 0.49 [2] and large-scale genome-wide association studies (GWAS) have been successful in identifying common variants associated with alcohol consumption and AUD related disorders. Together, common risk variants can account for up to 7-12% of the variance in AUD and related disorders [3–5].

Although pinpointing the biological effects of variants identified from GWAS of complex disorders is challenging [6], some of the results from studies of alcohol-related disorders represent an exception to this general pattern. The alcohol dehydrogenase 1B (*ADH1B*) gene encodes a subunit of the primary ethanol metabolizing enzyme. Across multiple GWAS of subjects of European descent, coding variant rs1229984 (Arg48His) is significantly associated with AUD [4, 7] or problem alcohol use[3, 5]. In subjects of African descent, rs2066702 (Arg370Cys) is significantly associated with AUD[7]. In both cases, the variant allele is associated with reduced AUD risk. The mechanism of this protective effect is not yet understood, but protein products encoded from either variant allele (containing His at position 48 or Cys at position 370) have a 70-80 times higher turnover rate for ethanol than protein products with Arg at positions 48 or 370[8]. The importance of these *ADH1B* coding variants suggests that analysis of coding variants identified by exome sequencing studies, most of which are rare, may identify additional genes contributing to risk and potentially reveal the mechanism of these effects.

GWAS assessing the role of common variation in complex traits require large samples to have power to detect the small allelic effects across a highly polygenic set of risk loci observed and have achieved these sample sizes through cost-effective microarray-based genotyping and imputation. Studies of rare coding variation in the exome also require large samples to have power to detect the effects of very low frequency alleles[9], even when aggregated over a biological system or biochemical pathway[10], and samples in excess of 100,000 subjects are needed to detect these effects aggregated over a single gene or individually, even in traits with high heritability of ∼ 0.8 like schizophrenia[11]. Although sequencing costs are falling, exome studies of many traits, including AUD and alcohol related disorders, remain underpowered. The two largest published exome studies of ascertained alcohol dependence or alcohol use disorder included approximately 10,000[12] and 25,000[13] subjects.

Biobank collections such as the UK Biobank (UKB), an ongoing survey of over 500,000 UK resident participants, represent one way to address the need for large samples to improve genetic discovery for other complex traits, including alcohol-related phenotypes. The UKB includes diverse measures of lifestyle, dietary habits, general and mental health, and health records, as well as microarray genotype, exome sequencing and whole genome sequencing data[14]. However, clinical diagnoses of AUD using structured interviews for Diagnostic and Statistical Manual of Mental Disorders (DSM) or International Classification of Disease (ICD) criteria are not feasible in biobank collections due to the need to collect a broad range of health and lifestyle data.

In such situations, screening instruments such as the Alcohol Use Disorders Identification Test (AUDIT) can be used[15]. The AUDIT is a screening questionnaire designed to identify hazardous alcohol use that consists of 10 items that produce a quantitative measurement from 0-40 across two subscales of alcohol consumption (AUDIT-C) and problematic use (AUDIT-P). Previous work has demonstrated that AUDIT-C shows strong genetic correlation with other consumption phenotypes such as drinks per week[16, 17], but only a moderate genetic correlation with DSM diagnosis of AUD. In contrast, there is a strong genetic correlation between AUDIT-P and DSM diagnosis of AUD, as well as other major psychiatric disorders such as schizophrenia and major depressive disorder, suggesting that AUDIT-P is useful for studies of alcohol related psychopathology when AUD diagnosis is unavailable because its use can increase statistical power[3, 4].

While the use of screening questionnaires such as the AUDIT provides a proxy measure of clinical diagnosis feasible for collection in a population sample, biobanks do not necessarily have consistent measurements for all variables across all subjects and may show extensive non- random block-wise missingness in their phenotypic and survey data which limits the effective sample size for genetic association studies[18]. This is a limitation for both common and rare variant studies, as a recent exome-wide analysis of the AUDIT using the 200k exome release of the UK Biobank failed to identify any significant genes or variants associated with heavy drinking or problem drinking[19]. This is likely attributable to inadequate sample size and power for rare variant association testing because only ∼30% of the 500k UK Biobank participants have completed the AUDIT questionnaire, and only ∼50k of these subjects are included in the 200k release.

One way to address the issue of inadequate power and effective sample sizes for rare variant studies is to utilize the available information through machine learning (ML) to predict the phenotype of interest in subjects for whom it is missing. We have previously developed an application of the Group LASSO called the Missingness Adapted Group-wise Informed Clustered LASSO (MAGIC-LASSO) which predicts unmeasured quantitative outcomes such as AUDIT with high phenotypic and genotypic accuracy in the full UK Biobank sample[18]. Another way to potentially increase the statistical power for rare variant identification is to incorporate functional genomics information as *a priori* weights.

In addition to protein-coding variants, previous work shows that variants impacting non- coding functional elements involved with gene regulation such as promoters, enhancers, chromatin conformation, and DNA accessibility are enriched in GWAS associations[20, 21] and incorporating such functional information can increase power in variant identification in GWAS studies[22]. While functional annotations are critical to increase our understanding of intronic and intergenic variants associated in GWAS and WGS studies, WES studies also include meaningful numbers of non-protein coding variants which map to promoters, enhancers, UTRs, introns, and some intergenic regions due to the capture of regions adjacent to exon targeting probes. As such, functional information can also be useful in WES studies, particularly if annotations specific to the phenotype of interest, which can be identified via enrichment analysis, are used in the analysis. One potential approach to increase power and specificity for rare variant analyses is to leverage results from GWAS of common variants as the basis for empirical functional weights for downstream rare variant testing.

In addition to the strong evidence across complex traits that functional annotations show enrichment, the specific annotations and magnitude of enrichment vary across disorders. For example, FANTOM5 enhancers[23] show strong enrichment for immunological disorders such as Crohn’s Disease or Ulcerative Colitis, while H3K4me3 annotations (marking active promoters) from neuronal cells are significantly enriched for psychiatric disorders such as schizophrenia[24]. Furthermore, complex disorders show convergence between common and rare variant signals, suggesting that incorporating information from GWAS and functional genomics could refine and increase discovery power for rare variant testing[25–27].

In this study, we analyzed unrelated European subjects within the 200k exome release of the UK Biobank to conduct an interval-based rare variant analysis on AUDIT-P. Sequence Kernel Association Test (SKAT-O) analyses were performed using subjects for whom AUDIT-P is directly measured (N=51,357) and predicted (n=147,376) irrespective of measured or unmeasured. The two analyses were compared to evaluate whether the increase in effective sample size can improve discovery power for rare variant association testing in AUDIT-P. Additionally, we evaluated the impact of including disease-specific functional information as *a priori* weights to investigate whether inclusion of functional information can improve statistical power for rare variant association testing compared to default SKAT/SKAT-O weights.

To our knowledge, this is the largest available rare variant study of AUDIT-P to date.

We hypothesize that by increasing the effective sample size and including disease-specific functional information as *a priori* weights for the interval testing, we may be able to uncover rare variant signals associated with AUDIT-P that would be missed using only directly measured subjects and default SKAT variants weights that only consider allele frequency. These results are expected to complement the growing literature on common variant studies on alcohol-related phenotypes.

## Methods

### UK Biobank Cohort

This research has been conducted using the UK Biobank dataset (application 30782). The UK Biobank dataset is a large, population-based sample that includes more than 500,000 participants aged between 40 and 69 years [28]. A wide range of phenotypic measurements and biological samples have been assessed and collected for these participants at different centers located in the UK. The UK Biobank has obtained ethics approval from the North West Multi- Center Research Ethics Committee, as well as informed consent from all the participants. In addition to the exome sequencing described in more detail below, individuals in the UK Biobank were also genotyped using the Affymetrix UK BiLEVE or Affymetrix UK Biobank Axiom arrays and the genotypes were imputed to the Haplotype Reference Consortium reference panel using IMPUTE2 [29] with full details provided elsewhere [30].

### Phenotype description and prediction of AUDIT-P using MAGIC-LASSO

The AUDIT is a ten-item, screening questionnaire instrument for alcohol consumption and problems containing three questions that survey consumption (AUDIT-C), and seven that survey problematic alcohol use (AUDIT-P). The survey was completed as part of the Mental Health Questionnaire (MHQ) battery in a subset of 157,162 out of the full 500k UK Biobank participants.

The MAGIC-LASSO procedure, described previously[18], was applied to predict AUDIT scores in participants for whom the AUDIT questionnaire was not directly administered in the UK Biobank. In brief, the MAGIC-LASSO procedure is an adaptation of the Group Least Absolute Shrinkage and Selection Operator LASSO (Group-LASSO) ML method for penalized regression[31] that can predict variables in the presence of non-random, block-wise missingness. The procedure represents a new implementation of established ML algorithms and employs a regression-based solution that is suited for the penalization of categorical predictors which are prevalent in the UK Biobank. MAGIC-LASSO procedure involves 1) characterizing missingness, 2) filtering variables for general missingness and balance across training and target sets, 3) variable clustering based on missingness, 4) iterative Group-LASSO and variable selection within clusters, and 5) cross-cluster model building with variables prioritized by informativeness. The phenotypic correlation between measured and predicted Total, Consumption, and Problems scores were 0.64, 0.71, and 0.48, respectively, while the LDSC based[32] genetic correlation (r_G_) between observed AUDIT in measured subjects and predicted AUDIT in unmeasured subjects (who are completely independent) was 0.92, 0.86, and 0.79 for Total, Consumption, and Problems respectively, demonstrating the method has significant accuracy and utility. For the present analysis, we focused only on the Problems subscale.

Importantly, these correlations indicate that the MAGIC-LASSO predictions effectively recapture unmeasured phenotypic information and likely lie closely along the same genetic continuum as observed AUDIT-P. More information is provided elsewhere in the original publication[18].

### Whole exome-sequencing in the UK Biobank

The 200k exome release of the UK Biobank dataset, including exome sequencing for 200,643 participants, was used in this analysis. Exomes were captured using the IDT xGen Exome Research Panel v.1.0 with an average coverage of 20X at 95.6% of sites. The Original Quality Functional Equivalent (OQFE) Pipeline was used to map raw FASTQ files with BWA- MEM to the GRCh38 reference genome while retaining all other alignments. The OQFE CRAMs were then called using DeepVariant to generate per-sample gVCFs that were jointly called using GLnexus. The OQFE version of the Plink formatted exome files was then downloaded and utilized for all the analyses described in this study (field 23155). Samples were initially filtered to retain only unrelated subjects of British ancestry (n = 359,980) as in previous analyses[33]. This yielded 147,376 participants with exome data from the 200k exome release of whom 51,357 had AUDIT directly measured.

### Empirical functional weights from predicted AUDIT-P GWAS

GWAS of common variants associated with predicted AUDIT-P was used as the basis for empirical functional weights for downstream rare variant testing. GWAS was conducted using the BGENIE software (version 1.3)[30], as described previously[18]. Briefly, pre-GWAS filtering excluded markers with MAF < 0.5%, INFO < 0.8, and HWE p-value < 10^(-6).

Association analyses included age, sex, and the first 20 ancestry principal components as covariates. After filtering, the sample of independent European subjects used for GWAS was 359,980 with 117,559 and 242,421 subjects in the AUDIT measured and unmeasured sets, respectively.

Stratified LD Score Regression (LDSC)[34] was used to partition the heritability of predicted AUDIT-P GWAS described above[18] into functional annotation classes while accounting for the overlap between different functional classes using the overlap-annot flag. In addition to the baseline functional annotation classes from LDSC V.2.2, four custom brain- specific functional annotation classes acquired from the psychENCODE Consortium[35] were used for stratified LDSC analysis which resulted in 87 annotation classes in total. The four custom annotations classes from the psychENCODE Consortium included 1) psychENCODE Enhancers, 2) h3k27ac markers in the prefrontal cortex, 3) h3k27ac markers in the temporal cortex, and 4) h3k27ac markers in the cerebellum. These datasets are publicly available and can be downloaded as part of the derived datasets from http://resourceT.psychencode.org/. The four annotations used are listed under “PsychENCODE enhancer list and H3K27ac peaks” in the “Derived Data” section and links to download each dataset are:

http://resource.psychencode.org/Datasets/Derived/DER-03a_hg19_PEC_enhancers.bed

http://resource.psychencode.org/Datasets/Derived/DER-05_PFC_H3K27ac_peaks.bed

http://resource.psychencode.org/Datasets/Derived/DER-06_TC_H3K27ac_peaks.bed

http://resource.psychencode.org/Datasets/Derived/DER-07_CBC_H3K27ac_peaks.bed

Empirical weights based on functional annotations were constructed as follows: 1) Each observed exome variant in the vcf was annotated for the presence or absence for each functional annotation class, 2) annotations were retained if significant by Akaike information criterion (AIC) in the stratified LDSC analysis, 3) the enrichment scores for significant annotation classes which represents the fold enrichment of that annotation class derived from stratified LDSC, were added up for each position in the exome. In cases where a variant did not fall within any of the significantly enriched annotation classes, the variant would not get up-weighted for the analysis and receive a weight of zero.

### Interval definitions

GENCODE V.39 was used as the basis for defining intervals to include the most comprehensive list of possible intervals in this analysis[36]. In addition to protein-coding genes, GENCODE includes gene models from multiple classes of noncoding RNA genes and other locus types, allowing for a more comprehensive examination of variation in expressed sequences. A comprehensive set of intervals across the genome including those between transcripts (e.g. intergenic intervals) was constructed. Due to overlapping protein coding transcripts, nested small RNAs, and long non-coding RNAs, constructed intervals were not necessarily independent. Although the current study used exome data, some intergenic intervals were observed to contain variants possibly due to off-target capture or annotation errors. All intervals regardless of class (protein coding, intergenic) were included in the analysis if they contained observed rare variation. Figure 1 shows the distribution of interval classes tested in our analyses. Of the 53,171 defined intervals, association testing was limited to intervals with at least 2 rare (MAF<0.01) variants. In total, 21,104 intervals were tested with most (n=17,968) being protein-coding genes, as expected.

**Figure 1:**
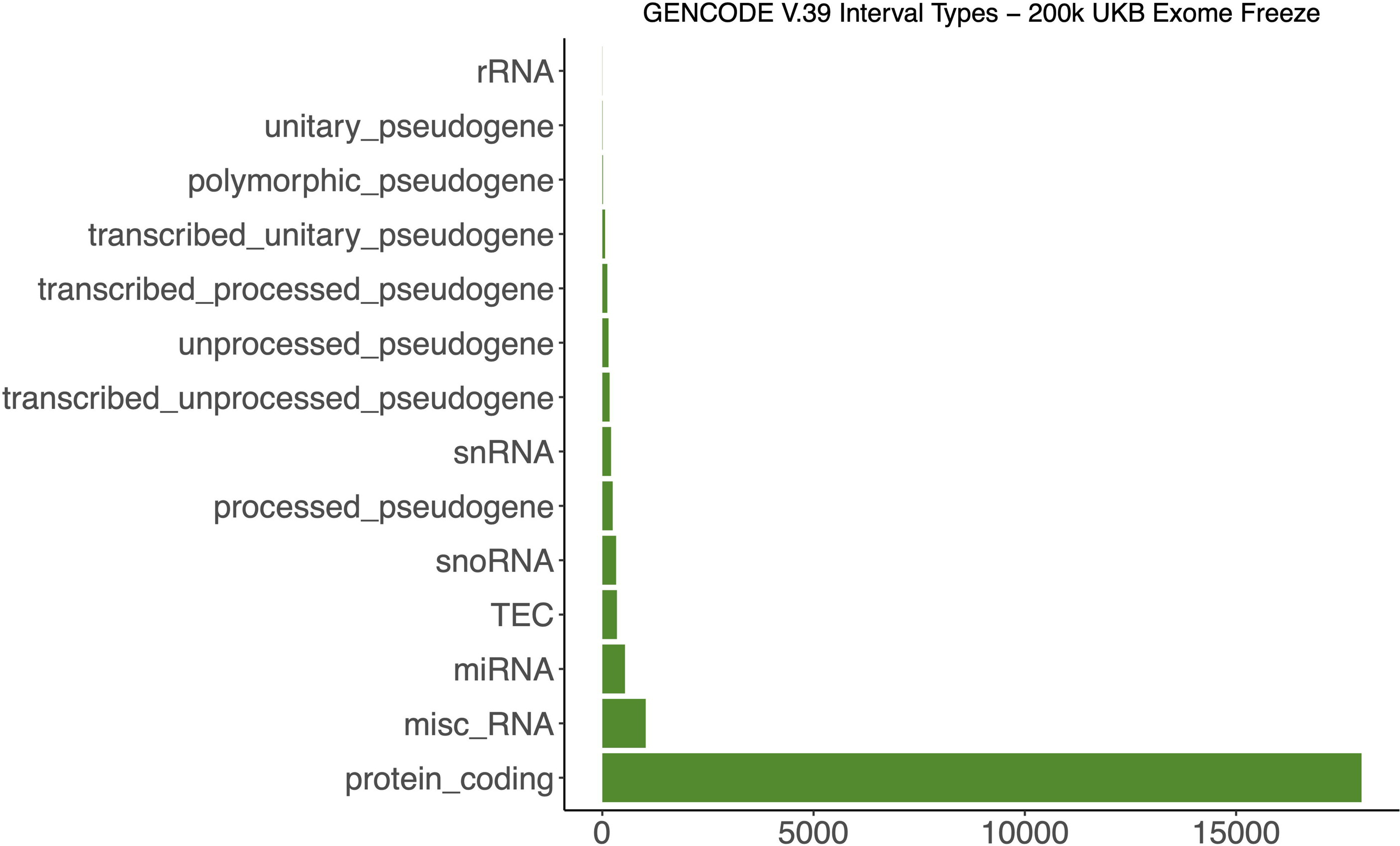
Distribution of the intervals tested in AUDIT-P rare variant analysis. Intervals were defined based on GENCODE V.39 definition of genic and non-genic intervals. In total, 21,105 intervals were tested across the exomes.

### Rare-variant interval-based association testing

Interval-based rare variant aggregate testing was performed using SKAT[37] and SKAT- O[38], as implemented in the SKAT package in R[39]. False discovery rate (FDR) analysis was performed and Q-values were generated for each test using the qvalue package in R[40]. FDR is more appropriate than family-wise error correction since gene intervals are not necessarily independent due to overlap and rare variant linkage disequilibrium (LD). In addition to FDR, we also reported corrected *p*-values using the conservative Bonferroni correction method. To assess inflation of *p*-values, we calculated the genomic inflation factor (lambda) for each set of tests, with a lambda value near 1 indicating no notable inflation[41].

Four sets of SKAT-O interval tests were carried out varying two conditions. First, SKAT- O tests were performed using default weights, which are based on allele frequencies, versus empirical functional weights constructed based on significant annotation classes from stratified LDSC analysis in subjects for whom AUDIT-P scores were directly measured (N=51,357). The goal was to evaluate whether *a priori* information from the enrichment of common variant GWAS data in combination with functional annotation information can improve statistical power for rare variant testing. Second, SKAT-O tests using the default versus empirical weights were performed in the full sample of 147,376 unrelated subjects with predicted AUDIT-P. The goal here was to investigate whether increase in effective sample size can increase our statistical power for rare variant testing. We hypothesized that using both a) empirical functional weights based on *a priori* functional annotation enrichments and b) increased effective sample size using predicted AUDIT-P would improve detection of intervals containing rare variants influencing the trait.

We opted to use SKAT-O for interval testing in this study. Burden tests are powerful when most variants in the interval are causal, with the effects in the same direction. Conversely, SKAT is more powerful when a fraction of the variants are noncausal, or have causal variant effects with mixed direction of effects. The SKAT-O approach maximizes power by adaptively using the data to combine burden and SKAT tests while maintaining the power for rare variant testing, making it an appropriate method when the direction of effects for causal variants are not known *a priori*.

## Results

### Partitioned heritability of functional annotations

Using LDSC, we estimated the heritability (h^2^) of the predicted AUDIT-P (N=359,980) to be h^2^ = 0.0647 (SE=0.0034), in close agreement with measured AUDIT-P in the UK Biobank (N=121,604)[3]. Partitioning the heritability of predicted AUDIT-P GWAS into functional categories resulted in 12 significant annotations after multiple testing corrections (Figure 2). Of these annotation classes, three were specific to the brain from the psychENCODE consortium (Table 1, full results in Supplemental Table 1) demonstrating the utility of using annotations beyond the available baseline set from LDSC.

**Figure 2:**
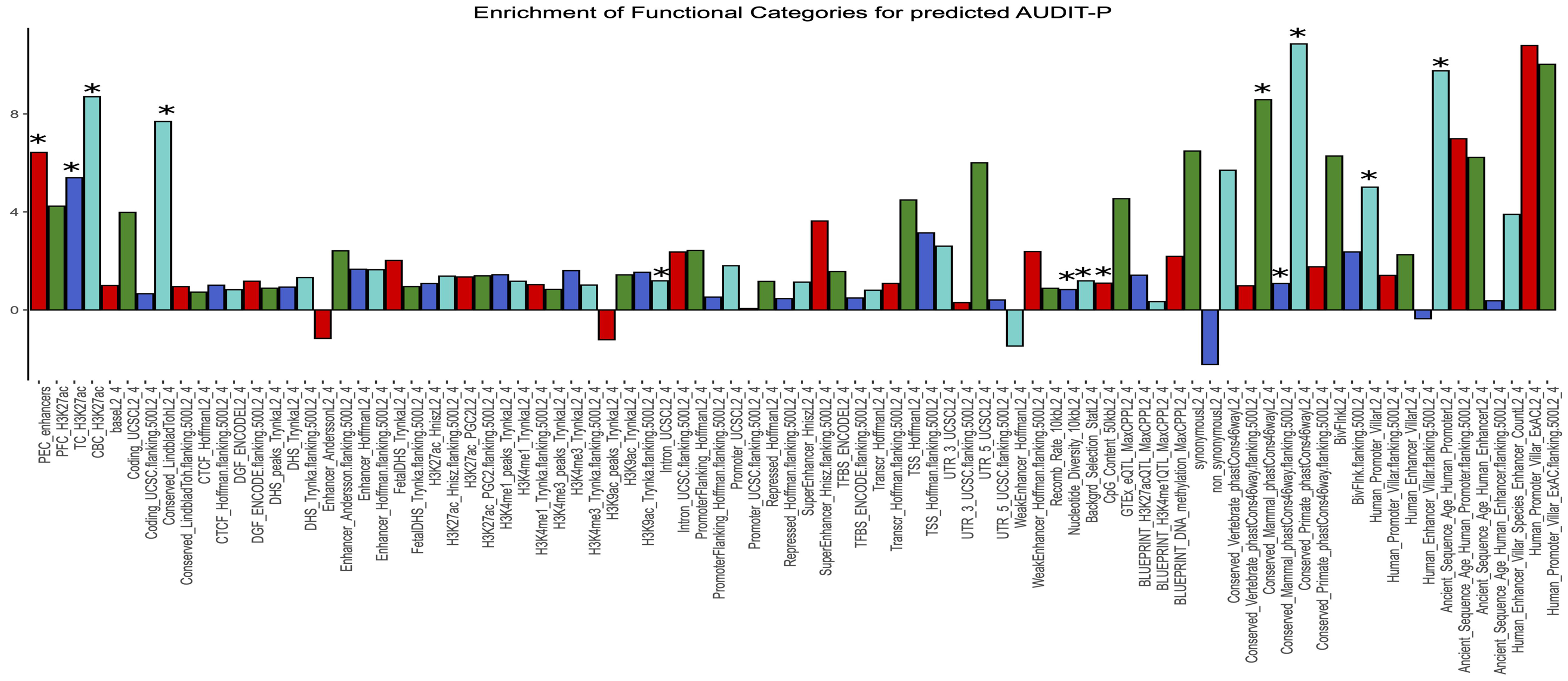
Partitioning the heritability of predicted AUDIT-P GWAS into functional categories using stratified LDSC. Asterisks represent significant enrichment classes after multiple testing correction. X-axis represents functional annotation classes and Y-axis shows the enrichment values.

**Table 1:**
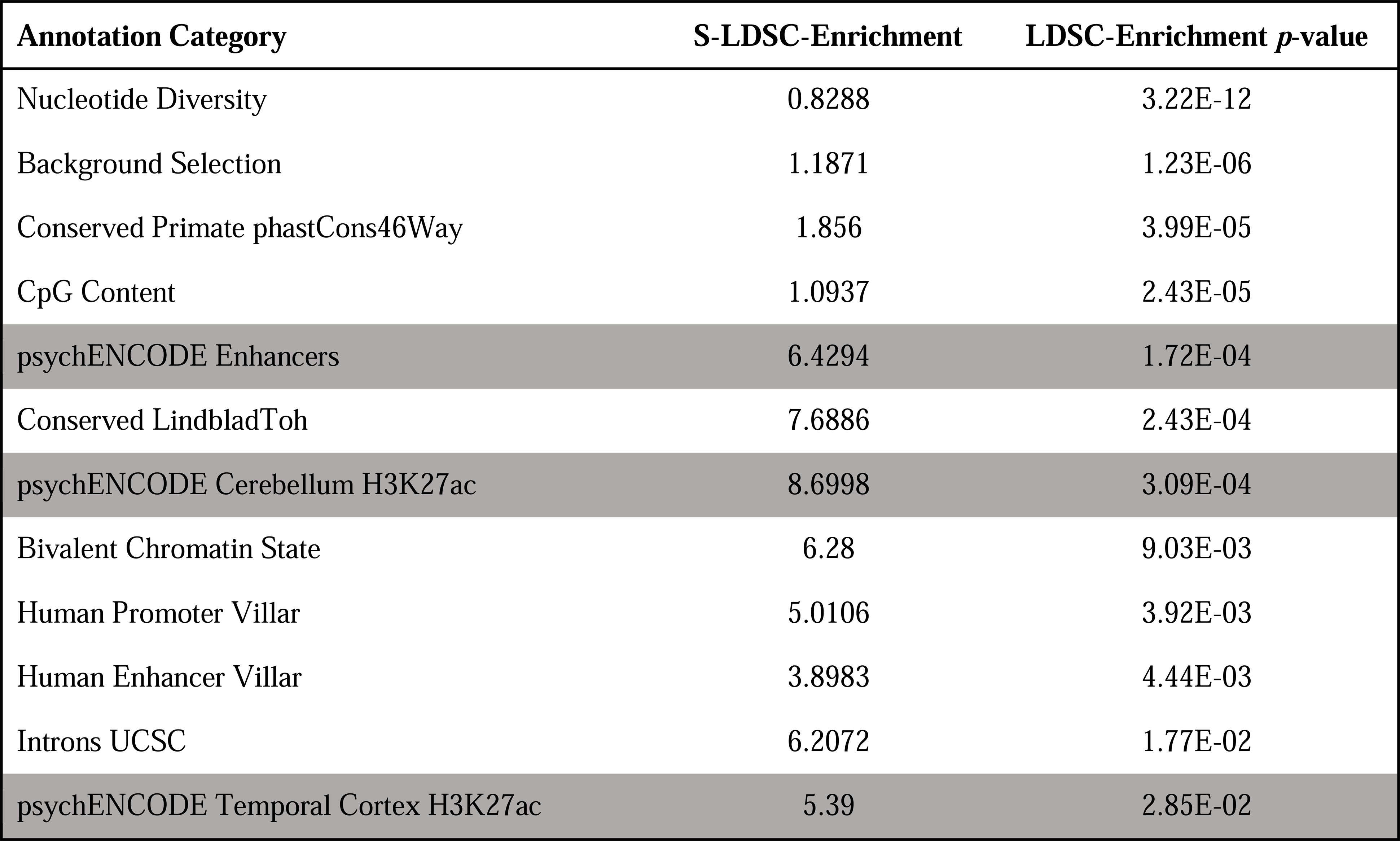
Significant annotation classes enriched in the heritability of predicted AUDIT-P after multiple testing correction. Grey categories are custom annotation classes.

Comparing the partitioned heritability results of predicted AUDIT-P with other complex traits reveals potentially important insights. For example, heritability of height GWAS[42] is significantly enriched in 36 functional classes. However, none of the brain-specific annotation classes from the psychENCODE consortium show significant enrichment for heritability of height (Supplementary Table 2). In contrast, heritability of schizophrenia GWAS[27] is significantly enriched in 21 annotation classes, which includes all four brain-specific annotation classes from the psychENCODE consortium (Supplementary Table 3). Finally, heritability of drinks per week GWAS[17] is significantly enriched in 10 functional classes, none of which are specific to the brain (Supplementary Table 4). While both schizophrenia and drinks per week show genetic correlation with predicted AUDIT-P, the pattern and magnitude of enriched annotation are different. Together, these results suggest that while there are important differences in heritability enrichment of these complex traits, inclusion of brain-specific annotation classes for psychiatric traits such as schizophrenia or AUDIT-P can improve stratified LDSC analyses and provide better disorder-specific empirical functional weights for rare variant testing.

### Empirical functional weights for AUDIT-P

The majority of the variants (63.4% of 17,549,750 variants) did not fall into any significant annotation classes. Therefore, these variants were not informative for empirical weight construction and received a weight of zero regardless of frequency. This is in contrast to the default weighting of SKAT which assigns weight based on MAF regardless of functional information. However, 36.6% of variants fell into at least one of the annotation classes. The resulting quantitative score based on combining the 12 significantly enriched annotations ranged from 0 to 71.23. Figure 3 shows the distribution of the empirical functional weights generated for all observed variants by allele frequency bin. Although not intended *a priori*, low frequency variants were overrepresented for higher weights.

**Figure 3:**
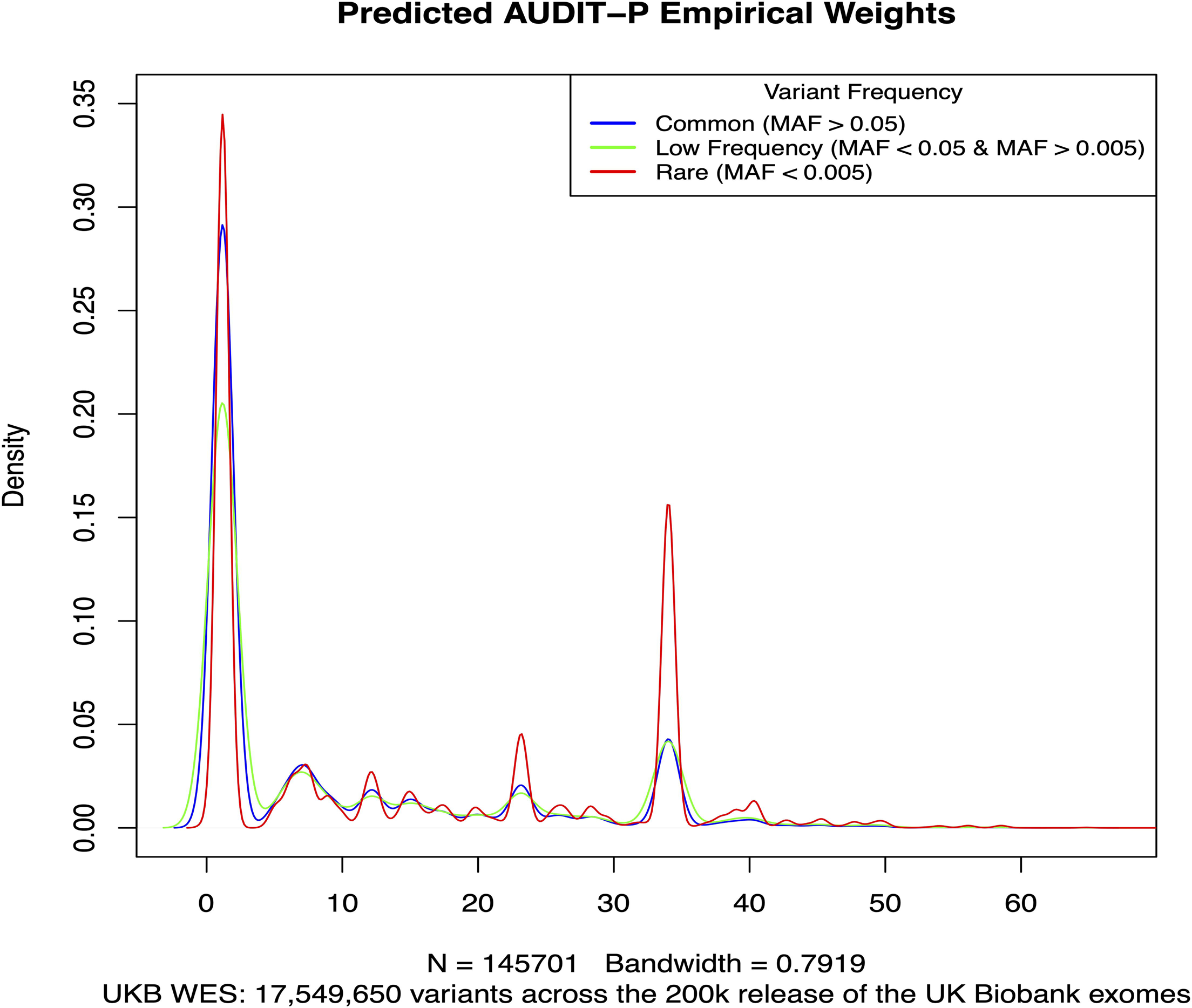
Distribution of empirical functional weights across the 200k UK Biobank exome dataset.

### Interval-based tests

Post filtering, interval-based analysis was performed using 17,59,650 qualifying variants that mapped onto 21,105 intervals. Figures 4 and 5 show SKAT-O interval-based results using subjects with directly measured (N=51,357) and predicted (measured and unmeasured, N=147,376) AUDIT-P, respectively. No interval was significantly associated with observed AUDIT-P, as previously observed, using the default (Figure 4A) or the empirical functional weights (Figure 4B). However, as shown in the QQ-plot (Figure 4C), empirical functional weights provide an improvement in statistical significance compared to default weights with no evidence of inflation when using empirical versus default weights. Empirical functional weights identify as many or more significant intervals than default weights across the FDR threshold bins (Table 2). Overall, there was little evidence for genomic inflation across all approaches, with lambdas using default and custom weights of 1.04, and 0.997 for SKAT-O, respectively. SKAT results for measured AUDIT-P are provided in the Supplementary Figure 1 and also did not show evidence of inflation with lambdas of 1.1 and 1.06 for default and empirical functional weights, respectively.

**Figure 4:**
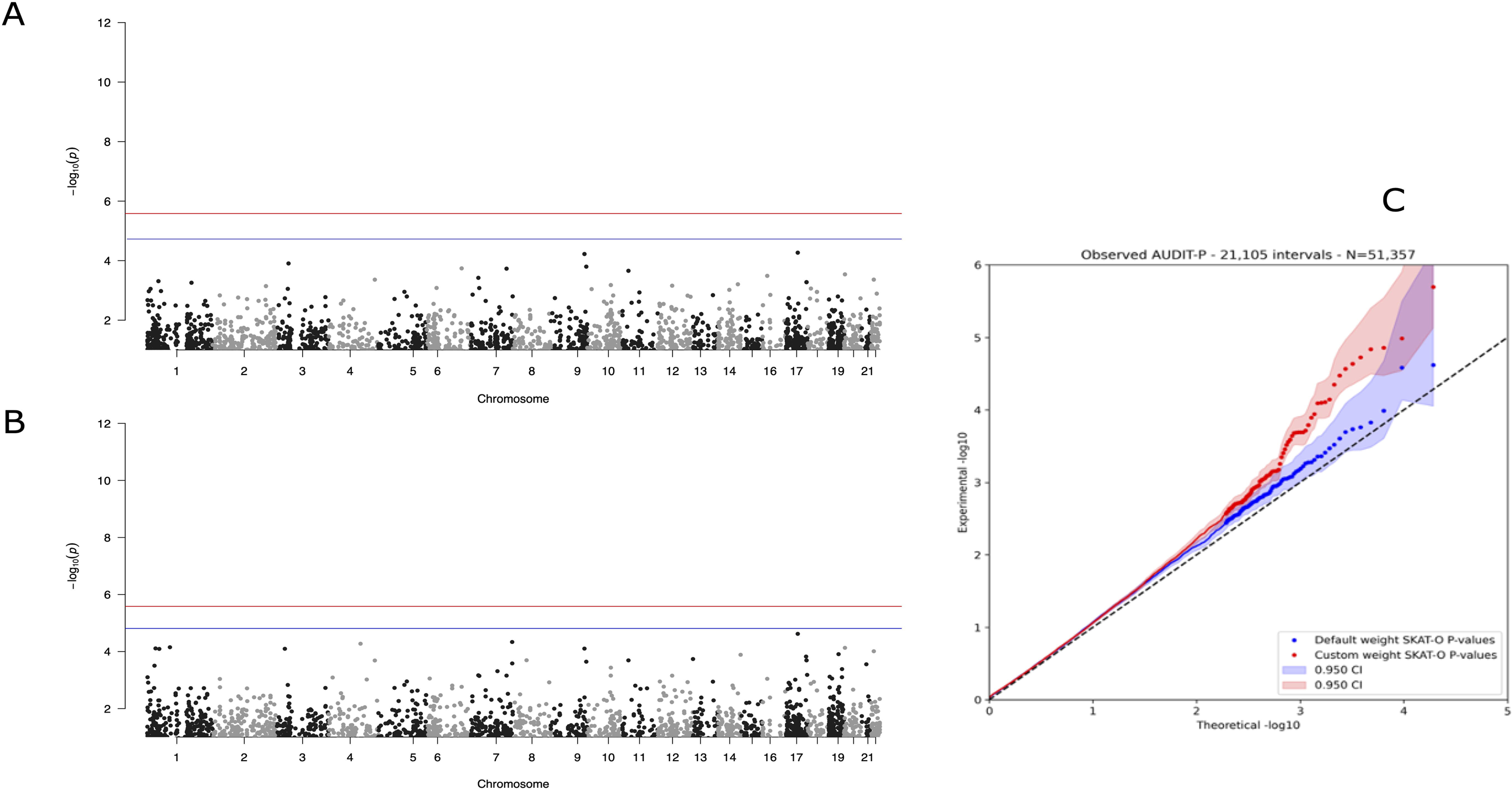
SKAT-O analysis on observed AUDIT-P using default (A) and empirical (B) weights. Q-Q plots for default and empirical weights are shown in panel C. For Manhattan plots, the red and blue lines represent conservative Bonferroni correction and FDR thresholds of 5%, respectively.

**Figure 5:**
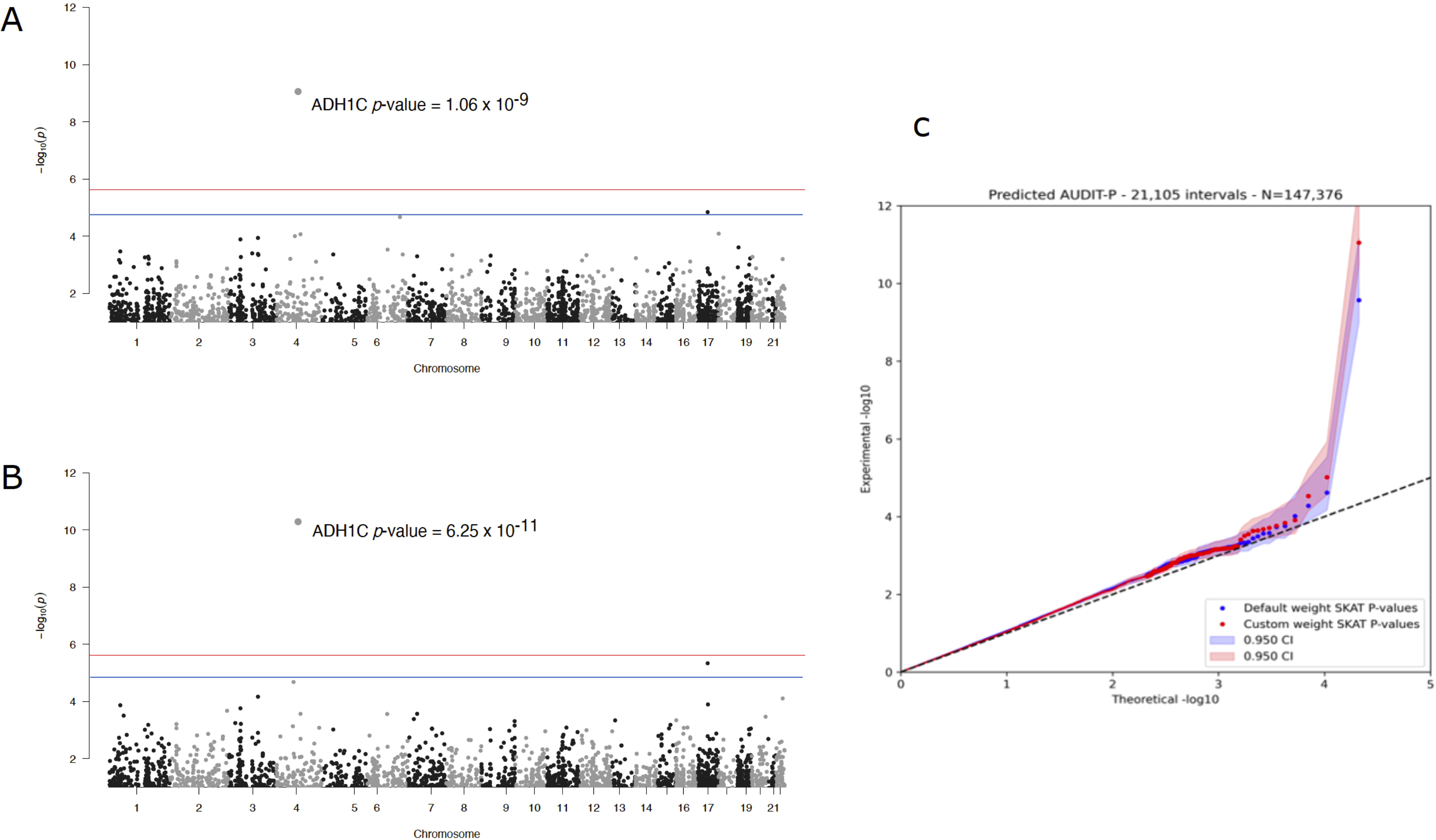
SKAT-O analysis on predicted AUDIT-P using default (A) and empirical (B) weights. Q-Q plots for default and empirical weights are shown in panel C. For Manhattan plots, the red and blue lines represent conservative Bonferroni correction and FDR thresholds of 5%, respectively.

**Table 2:**
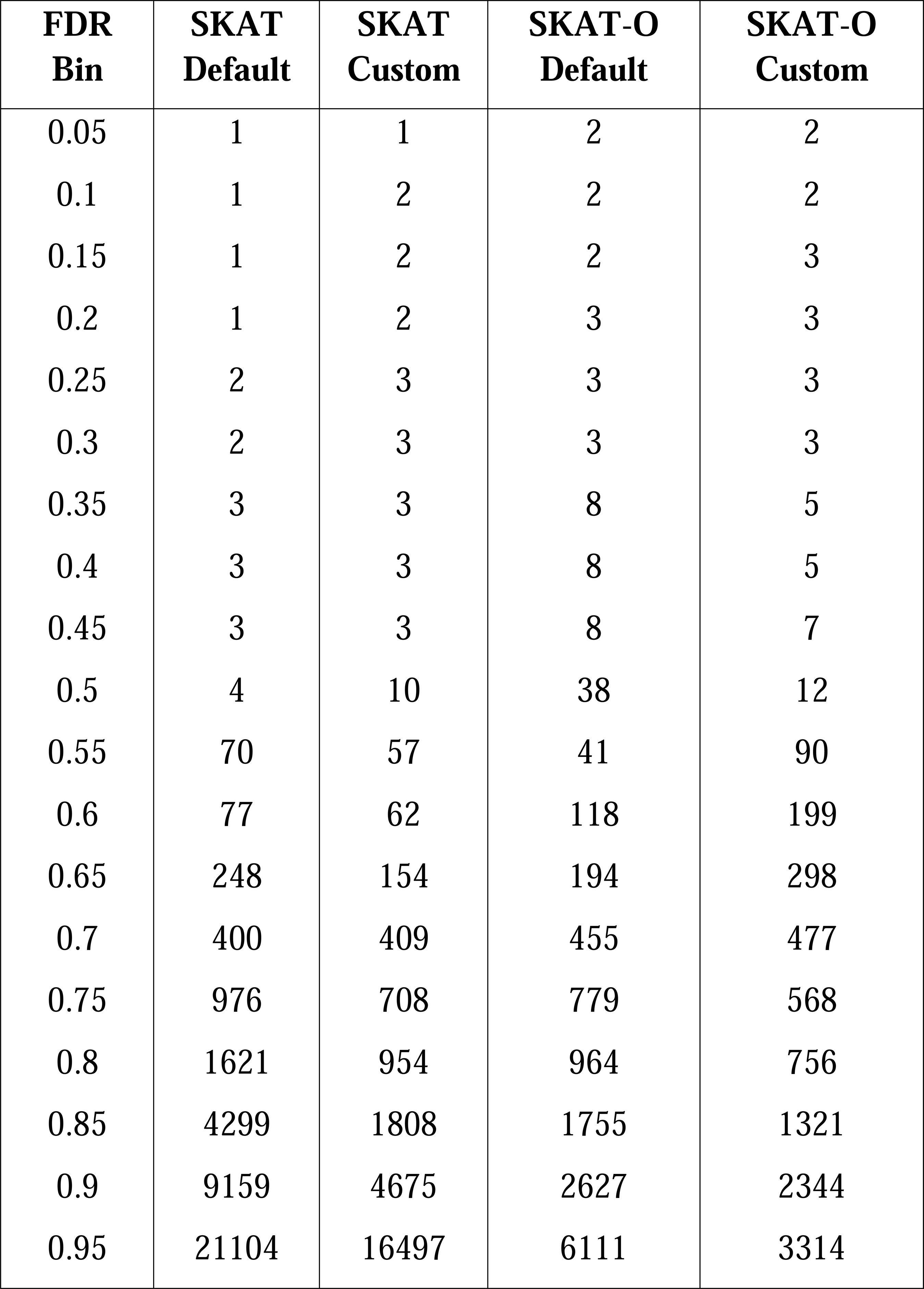
Count of intervals at or below given FDR thresholds for all four analysis sets, SKAT with default and custom weights, and SKAT-O with default and custom weights.

In contrast to using directly measured AUDIT-P, we observed significant associations with *ADH1C* and *THRA* gene intervals for predicted AUDIT-P when using both default (Figure 5A) and empirical functional weights (Figure 5B). A two-fold increase in statistical significance was observed when using empirical functional weights compared to default weights (Figure 5C), with the most significant association found with predicted AUDIT-P and empirical weights in the *ADH1C* gene after conservative Bonferroni correction (SKAT-O P*_Default_*= 1.06 x 10^-9^ and P*_Empirical Weights_* = 6.25 x 10^-11^). SKAT results for predicted AUDIT-P are provided in the Supplementary Figure 2.

*The ADH1C* gene interval which encodes class I alcohol dehydrogenase gamma subunit has been previously implicated in alcohol-related phenotypes. Of the 670 variants in the *ADH1C*, 43 had MAF < 0.01 and were tested in the analysis. In contrast, the *THRA* gene interval encodes one of the receptors for thyroid hormone and mutations in this gene are associated with intellectual disability and reduction in brain size[43]. Of the 1,245 markers in *THRA*, 54 had MAF < 0.01 and were tested in the analysis.

## Discussion

In this study, we sought to perform an exome-wide rare variant analysis on potential alcohol problems using the AUDIT-P subscale in the UK Biobank sample. While common variant GWAS of alcohol consumption have successfully identified many common variant loci, studies of PAU and AUD have been less successful[3, 5] and large-scale rare variant analyses have yet to identify robustly associated variants, genes, or intervals[19]. To address this, we used ML phenotype prediction in the UK Biobank to increase the effective sample size for AUDIT-P for rare variant association testing to perform the largest exome-wide analysis of AUDIT-P to date. Additionally, while default SKAT weights are based on the frequency of the variants and thus not disorder specific, we incorporated evidence from common variant GWAS data and functional information as *a priori* weights to provide disorder specific weights for rare variant interval testing. Our findings show that in addition to common risk variation[5], rare risk variation in the *ADH1C* gene is also associated with AUDIT-P in the UK Biobank exome dataset, thus providing evidence for the involvement of rare risk variation in the genetic architecture of AUDIT-P at the population level.

In our first-pass analysis, we used the subset of the UK Biobank cohort with measured AUDIT-P (N=51,357) and found no significant association with rare variation using default or empirical functional weights. While we were able to demonstrate that including disorder-specific empirical functional weights appears to increase the statistical power (Figure 4C), we note that these *a priori* weights were insufficient to rescue an underpowered phenotype for rare variant association testing. Thus, larger sample sizes are required to adequately increase statistical power for rare variant identification in AUDIT-P. Conversely, by leveraging ML phenotype prediction to increase the effective sample size for rare variant association testing, we show that *ADH1C* and *THRA* genes are robustly associated with predicted AUDIT-P in the UK Biobank at rare variation level. Furthermore, we were able to demonstrate that inclusion of disorder-specific empirical functional weights generated from common variant GWAS data and functional genomics information, can also refine and increase the statistical significance for rare variation testing (Figure 5C). These findings highlight the successful leveraging of ML to substantially increase effective sample size to improve statistical power, and disorder-specific empirical functional weights to refine and increase the statistical significance in interval-based rare variant testing.

The protein products of the *ADH1A*, *ADH1B*, and *ADH1C* genes form homo-or hetero- dimers that convert ethanol to acetaldehyde and play important roles in hepatic alcohol clearance[44]. Common variants in *ADH1B* are robustly associated with both AUD diagnosis and AUDIT-P[3–5, 7, 16, 45, 46]. Therefore, our finding builds on previous common variant evidence for the involvement of variation in the *ADH* gene cluster in the genetic architecture of alcohol-related phenotypes by demonstrating that rare variation in *ADH1C* is also associated with alcohol problems as indexed by AUDIT-P. Similar to other psychiatric disorders such as schizophrenia[11], these results also suggest that there is a convergence between common and rare variant signals in the genetic architecture of AUDIT-P, and as sample sizes continue to increase, we can expect to identify more rare variant signals in previously implicated genes from common variant studies of alcohol-related phenotypes.

The *THRA* gene (encoding the thyroid hormone alpha receptor) has not been previously implicated in alcohol-related phenotypes. Thyroid hormone deficiency during pregnancy is a common cause of intellectual disability[47] as well as neurocognitive deficits, reduction in cerebellar volume and decreased white matter density in adults[43]. A recent study conducted in the UK Biobank has also demonstrated that increase in alcohol consumption is associated with decrease in global brain volume measurements as well as white matter microstructure[48]. Our finding suggests that rare variation in thyroid hormone receptors encoded by the *THRA* gene could also be involved in the development of potential alcohol problems.

Applying the MAGIC-LASSO to predict unmeasured AUDIT-P outcomes increased the sample size from 51,357 to 147,376, representing a 129% increase in effective sample size, after accounting for the phenotypic correlation between the observed and predicted AUDIT-P scores. Importantly, these represent AUDIT-P scores for 96,019 independent subjects with exome variant data who could otherwise not have been included in this analysis without ML predicted scores. While ML-predicted outcomes are not without error, we have demonstrated an approach for efficiently and reliably predicting missing outcomes to maximally leverage measured exome data for rare variant study of a phenotype such as AUDIT-P in the UK Biobank.

Predicting the likely functional impact of variation in the genome is challenging, as it requires taking appropriate account of the different kinds and levels of prior evidence of the likely function for every position. Previous analyses[20, 21] have shown that joint modeling of available functional information can improve power to detect putative causal variants in common variant studies across complex traits. In this analysis, we hypothesized that based on the evidence for the convergence of common and rare variant signals in the genetic architecture of complex psychiatric traits, similar approaches can be utilized to also improve signal detection in rare variant studies of complex psychiatric traits.

To achieve this, we first analyzed predicted AUDIT-P GWAS data by partitioning its heritability into functional sequence classes using stratified LDSC which accounts for the overlap among different functional classes. Enrichment values from statistically significant classes were then used to determine the overall variant weight at each position in the exome, by simultaneously accounting for the SNP-based heritability of common variant signals conferring risk to AUDIT-P, as well variant membership in each of the significantly enriched functional classes. We hypothesized that these disorder-specific weights would outperform default SKAT weights which assigns the weights to the variants based on the MAF without taking *a priori* functional information into consideration.

Our findings show that by leveraging common variant GWAS results specific to the phenotype to be used in a rare variant investigation as well as functional information, we can refine and increase the statistical significance. In this case, the application resulted in discovering two independent gene intervals containing rare variants influencing alcohol problems as measured by AUDIT-P. Additionally, of the four custom annotation sources acquired from the psychENCODE Consortium, three were significantly enriched for the heritability of AUDIT-P. This finding demonstrates that by going beyond default annotation classes from LDSC and including custom annotation classes such as enhancer and acetylation marks from the psychENCODE Consortium, we can provide more specific weights that can further increase detection power. While these findings can be seen as a proof of principle for the utility of including functional information as *a priori* weights in rare variant testing, given that most of the functional variation in the genome lies outside of the coding regions of the genome[49], future work could use whole-genome sequencing data to further explore the utility and feasibility of using empirical functional weights in non-coding regions of the genome. Furthermore, while AUDIT-P is moderately heritable, empirical functional weights derived from more heritable disorders such as schizophrenia[27] may show better predictive power for rare variant identification than AUDIT-P. Therefore, we could also explore the utility of empirical functional weights in more heritable disorders.

The analyses presented here should be interpreted in the context of some limitations.

First, there are currently no large-scale samples for rare variant studies of alcohol-related phenotypes which motivated us to use ML to predict AUDIT-P in the full 200k exome release of the UK Biobank for rare variant analysis. While we note that our predicted AUDIT-P shows strong genetic correlation with measured AUDIT-P in the UK Biobank (∼0.91), replication of the results presented in this study in other adequately powered cohorts is important. Second, due to sample size and power limitations, the current analysis utilized interval-based rare variant testing and thus, we did not conduct single marker tests. As larger sample sizes become available, it is important to extend these analyses to also perform single marker tests on AUDIT-P. Third, while our empirical functional weighting scheme shows improvement over the default SKAT weights, we note that future work could focus on refining the weights and applying it to WGS data where most variants are not limited to the exome. Fourth, the current analysis was limited to the European subset of the UK Biobank exome data release. As larger, more ancestrally diverse samples with adequate power for rare variant testing become available, future studies should conduct rare variant association testing on AUDIT-P in under-represented and diverse populations. Finally, the informativeness of the empirical weights is a function of the annotation sources and enrichment analyses. Many potential biological annotations including those that are tissue specific were not available or included. As annotation sources grow in number, tissue specificity, and underlying sample size, the utility of the empirical weight may improve.

In conclusion, in this study, we show that rare variation in the *ADH1C* and *THRA* genes is significantly associated with AUDIT-P in the UK Biobank. These results suggest that leveraging ML phenotype prediction and empirical functional weights can help increase effective sample size and subsequent discovery power for rare variant association testing in underpowered phenotypes such as AUDIT-P. As sample sizes increase, future directions of this work include improvement of the empirical functional weights and carrying out these analyses in the non- coding regions of the genome using whole-genome sequencing data.

## Acknowledgements

This research was supported by P50AA022537 (MA, AEG, MFH, THN, KSK, SAB, REP, BPR, BTW, PI Miles), R01MH114593 (MA, SAB, BPR, BTW, PI Riley), K25AA030072 (THN), and T32MH020030 (AEG, PI Neale). This research has been conducted using the UK Biobank Resource application number 30782. UK Biobank is generously supported by its founding funders the Wellcome Trust and UK Medical Research Council, as well as the British Heart Foundation, Cancer Research UK, Department of Health, Northwest Regional Development Agency and Scottish Government. The organization has over 150 dedicated members of staff, based in multiple locations across the UK. The UKB data utilized in this research is available to, “bona fide researchers for health-related research in the public interest,” through an application process accessible through the UKB website, https://www.ukbiobank.ac.uk/.

