## Supplementary figures and images for "Improving the discovery of rare variants associated with alcohol problems by leveraging machine learning phenotype prediction and functional information"

### Supplemental Figure 1

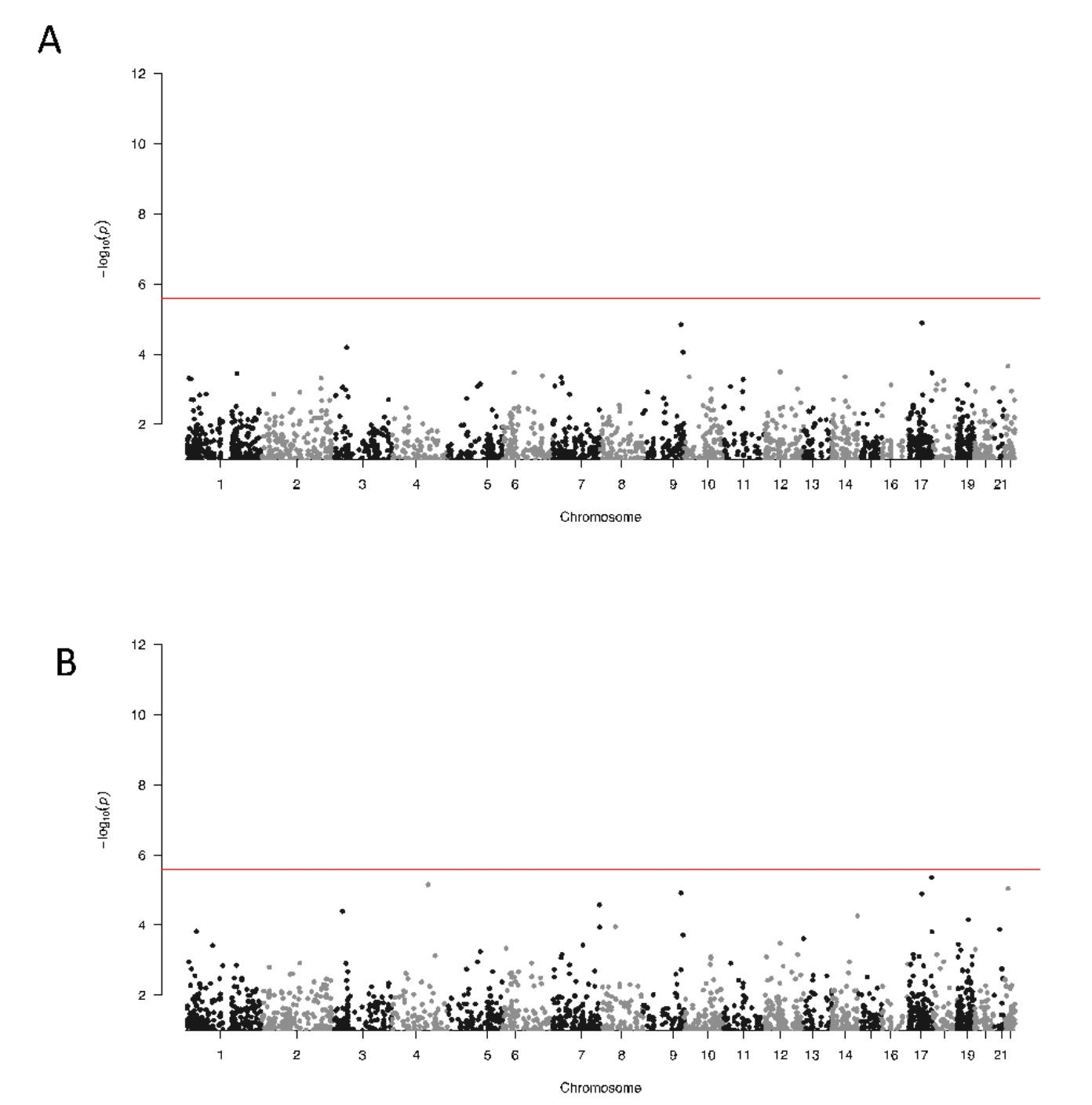

### Supplemental Figure 2

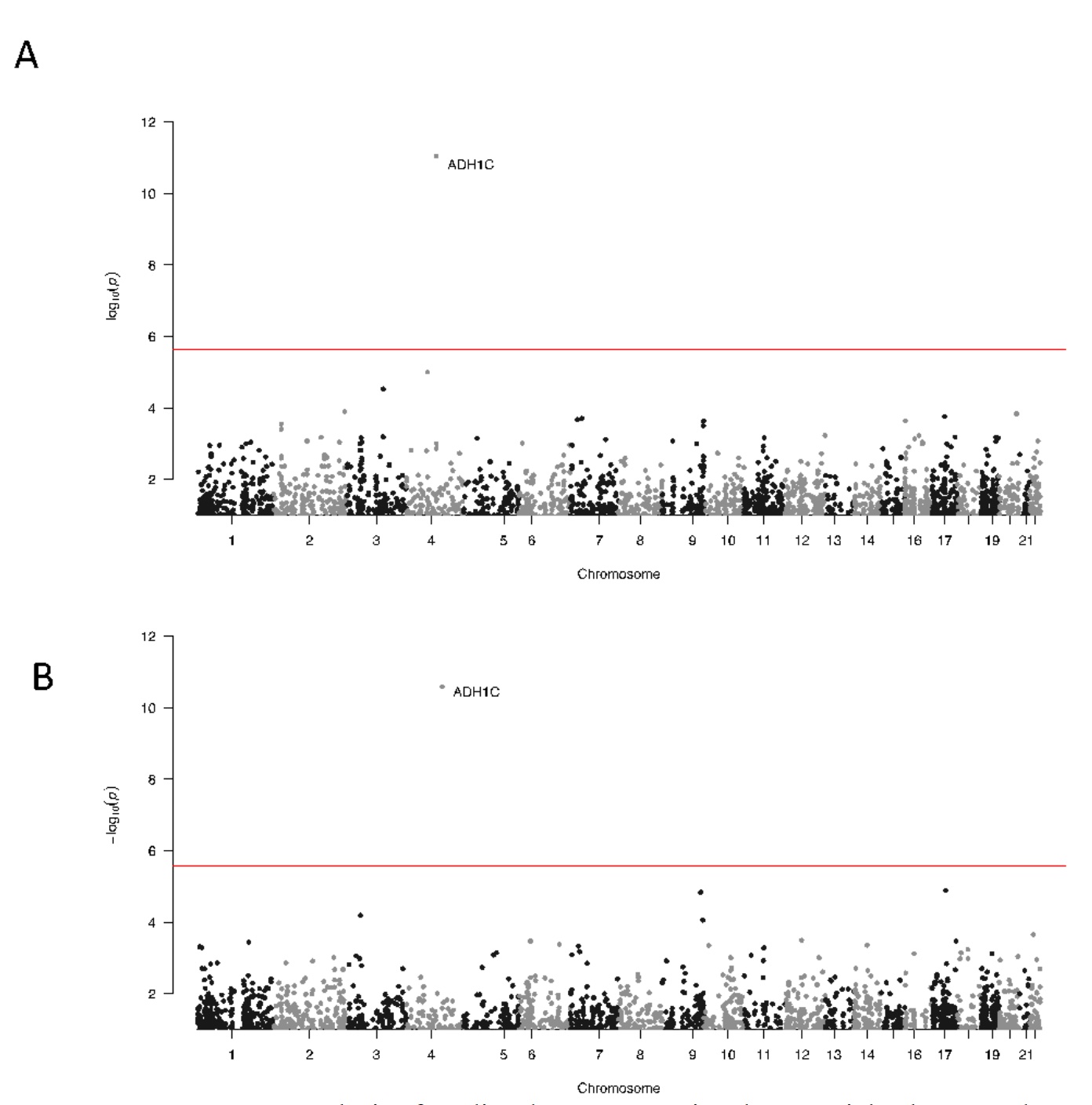
